# A pistillate-biased epidermal regulatory program underlies glandular trichome development in *Cannabis sativa*

**DOI:** 10.64898/2026.05.26.727882

**Authors:** Adrian S. Monthony, Julien Roy, Mohsen Niazian, Thomas Jarrin, Davoud Torkamaneh

## Abstract

Glandular trichomes are specialized epidermal structures on *Cannabis sativa* L. inflorescences that synthesize and store cannabinoids and terpenoids, making them central to the species’ economic and medicinal value. Although trichome development is a multistage genetic process, its regulatory basis in *C. sativa* remains poorly understood, particularly with respect to sexual dimorphism and sex plasticity. Here, we combined orthology-based gene discovery with transcriptomic profiling to investigate trichome development across four floral phenotypes: female flowers (FF), male flowers (MF), induced male flowers on XX plants (IMF), and induced female flowers on XY plants (IFF). Using trichome development-related genes from *Arabidopsis thaliana*, a model for unicellular non-glandular trichome development, and *Solanum lycopersicum*, a model for multicellular glandular trichomes, we identified and mapped 53 candidate *C. sativa* trichome development regulator genes (CsTDRGs). The CsTDRG set did not support a simple Arabidopsis– or tomato-like model, but instead included Arabidopsis-like epidermal fate components, including MBW-related regulators and *GL2*, alongside tomato-like multicellular trichome regulators, including *MIXTA-like, HD-ZIP IV, WOX, MTR, GRAS,* and hormone-associated candidates. RNA-seq analysis showed that CsTDRG expression was more strongly associated with floral phenotype than chromosomal sex. Genes with preferential expression in pistillate tissues were *CsTT8*, *CsMYC1/GL3*, *CsGL2, CsYABBY4* and *CsMIXTA-like1/MYB106*, whereas *CsMTR1*, *CsGRAS9*, and *CsCKX3* were upregulated in male (MF and IMF) flower. These findings suggest that sexually dimorphic trichome development in *C. sativa* reflects differential regulation of a shared developmental toolkit that combines conserved epidermal fate components with multicellular and glandular trichome regulatory modules.

## Introduction

*Cannabis sativa L.* (Cannabaceae) is an economically and medicinally important plant species known for its rich profile of secondary metabolites. This diploid (2n=20), typically dioecious species produces separate male and female individuals and can exhibit sexual plasticity, whereby under certain hormonal or chemical treatments (e.g., silver thiosulfate, ethephon, or gibberellic acid), genetically female plants can be induced to develop male flowers, and vice versa (Garcia-de Heer et al., 2025; Monthony et al., 2024, 2026). Leveraging sexual plasticity in *C. sativa* has enabled the production of feminized seeds (Flajšman et al., 2021), and provides a powerful experimental system for dissecting the molecular regulation of sex-related traits.

One sex-related trait of particular interest in *C. sativa* is trichome development. Cannabis produces multiple trichome morphotypes, including the glandular capitate-stalked, capitate-sessile, and bulbous forms, as well as several non-glandular types such as cystolith-bearing and simple non-glandular trichomes (Andre et al., 2016; Tanney et al., 2021). Among these, capitate-stalked glandular trichomes are the dominant trichome type on mature pistillate inflorescences and are most strongly associated with the biosynthesis and storage of specialized metabolites, including cannabinoids and terpenes (Livingston et al., 2020; Tanney et al., 2021; Figure 1). These compounds have attracted substantial interest because of their psychoactive properties, therapeutic potential, and commercial or industrial applications, including Δ9-tetrahydrocannabinol (THC), cannabidiol (CBD), and diverse terpenes (Mudge et al., 2018, 2019; Tetali, 2019). To date, more than 100 cannabinoids and over 150 terpenoids have been identified in cannabis (Andre et al., 2016; Hanuš et al., 2016). A similarly important role for glandular trichomes is seen in the related Cannabaceae species *Humulus lupulus* (hops), where they produce the α– and β-acids that contribute to beer bitterness (Carey et al., 2026). Together, the abundance and economic importance of trichome-derived specialized metabolites across the Cannabaceae underscore the need to expand current knowledge of trichome development-related genes (TDRGs) in this still underexplored group.

**Figure 1.**
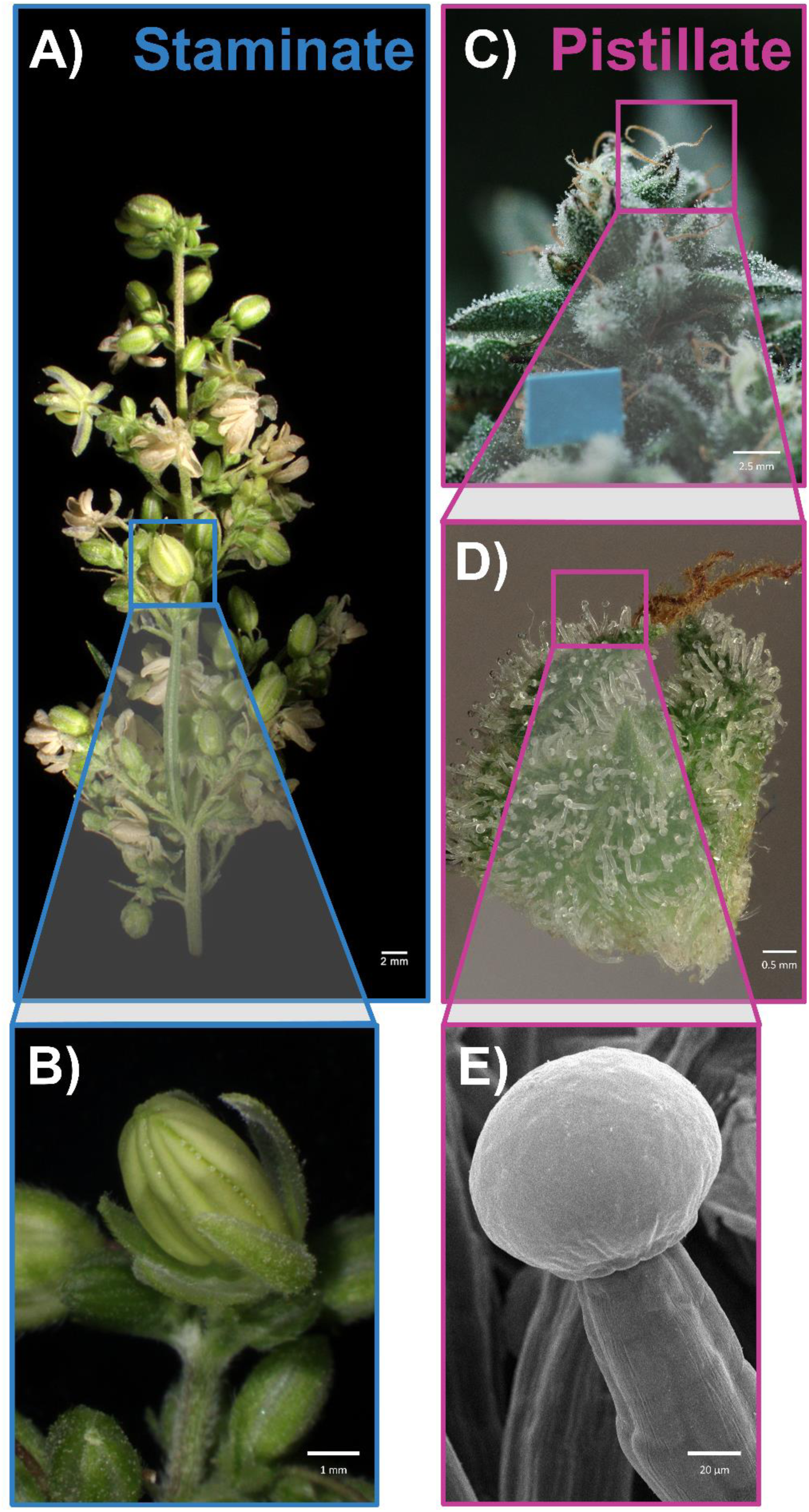
Sexual dimorphism and glandular trichome morphology in *C. sativa* floral tissues. (A) Representative staminate inflorescence showing male reproductive structures. (B) Magnified view of a staminate flower bud showing anther-bearing reproductive tissue. (C) Representative pistillate inflorescence showing female reproductive tissues and trichome-bearing floral surfaces. (D) Magnified view of pistillate floral bracts illustrating the dense distribution of glandular trichomes. (E) Scanning electron microscopy (SEM) image highlighting the morphology of a capitate-stalked glandular trichome. Scale bars: A, 2 mm; B, 1 mm; C, 2.5 mm; D, 0.5 mm; E, 20 μm.

The genetic mechanism underpinning glandular trichome development in *C. sativa* remains under-explored due to years of restriction on its cultivation and research. However, given the widespread prevalence of trichomes in many plant species, the process has been well-studied in model species such as *A. thaliana* and *Solanum lycopersicum* (Marks, 1997; Zhao et al., 2025). These species present particularly important models for trichome development, highlighting the pathway for single cell non-glandular trichome development (*A. thaliana*) and multicellular glandular and non-glandular trichome development (*S. lycopersicum)*. Trichome initiation in both species requires a genetically mediated change in epidermal cell fate: In unicellular, non-glandular trichomes such as those found in *A. thaliana* this process involves endoreduplication-driven cell expansion and the elaboration of a branched unicellular structure (Ilgenfritz et al., 2003; Marks, 1997). Multicellular glandular trichomes develop through a more complex and less well-understood process whereby epidermal cells proceed through coordinated rounds of cell division, cellular expansion, and differentiation into specialized secretory and supporting cell types. In some types, such as the glandular trichomes of *S. lycopersicum*, this process culminates in the formation of a subcuticular storage cavity where secondary metabolites accumulate (Hancock et al., 2024; Livingston et al., 2020; Raman et al., 2017).

In *A. thaliana*, trichome fate specification is regulated by a well-characterized MYB–bHLH–WD40 transcriptional module involving *GLABRA1* (*GL1*; R2R3-MYB), the bHLH proteins *GLABRA3* (*GL3*) and *ENHANCER OF GLABRA3* (*EGL3*), and the WD40-repeat protein *TRANSPARENT TESTA GLABRA1* (*TTG1*). This complex activates downstream regulators, including the HD-Zip IV transcription factor *GLABRA2* (*GL2*), which is required for normal trichome differentiation and epidermal patterning (Wang et al., 2010). Importantly, this regulatory architecture is not restricted to trichome initiation. Related TTG1-dependent MYB–bHLH–WD40 complexes also regulate root hair patterning, seed coat development, and flavonoid metabolism, with different MYB and bHLH partners conferring tissue– and pathway-specific outputs. For example, *TRANSPARENT TESTA 8* (*TT8*), a bHLH factor best known for its roles in anthocyanin and proanthocyanidin biosynthesis, also contributes to marginal trichome development in *A. thaliana*, revealing partial functional overlap among *TT8*, *GL3*, and *EGL3* within the broader *TTG1* network (Maes et al., 2008). More recently, jasmonate and ethylene signaling were shown to converge on WD40–bHLH–MYB complexes through *JAZ* repressors and *EIN3/EIL1* transcription factors, modulating anthocyanin accumulation, trichome formation, and defense responses (Song et al., 2022). This shared regulatory logic suggests that genes identified primarily in the context of specialized metabolite pathways, such as *TT8*, may nonetheless participate in the broader transcriptional networks underlying epidermal cell fate decisions, including trichome development.

In *S. lycopersicum*, multicellular trichome development is regulated by a partially distinct network centered on the HD-Zip IV transcription factor *WOOLLY* (*Wo*), which functions as a major regulator of trichome initiation, fate specification, and morphogenesis. Tomato produces seven major trichome types, including long multicellular type I trichomes, shorter non-glandular types II, III, and V, and glandular types I, IV, VI, and VII, with type VI trichomes serving as major sites of volatile terpene biosynthesis (Zhao et al., 2025). Although Wo is not a direct orthologous replacement for the Arabidopsis MBW complex, it occupies a broadly analogous epidermal fate-regulatory position to Arabidopsis HD-Zip IV factors such as *GLABRA2* (*GL2*). Additional regulators, including the C2H2 zinc-finger protein *Hair* (*H*), interact with *Wo* and are required for type-I trichome formation (Chang et al., 2018). Recent work has shown that *Wo* acts in a dosage-dependent manner, with higher *Wo* levels favouring digitate trichome fate through *SlWox3b/MX1* and lower Wo activity permitting peltate glandular trichome development through *LEAFLESS* (*LFS*) (Wu et al., 2023). Subsequent work further showed that a Wo protein gradient coordinates polarized cell division and basal cell expansion during multicellular trichome morphogenesis through *SlBRC2a* (Wu et al., 2024). In glandular trichomes, developmental regulation is also tightly integrated with specialized metabolism: Wo directly binds terpene synthase promoters and recruits the JA-responsive bHLH factor *SlMYC1* to activate terpene biosynthesis, while *SlJAZ2* represses this module in the absence of JA (Hua et al., 2021). Thus, tomato provides a complementary model to *A. thaliana* for identifying candidate regulators of multicellular and glandular trichome development in *C. sativa*, particularly where developmental regulation intersects with specialized metabolism.

The genetic regulation of trichome development in *C. sativa* remains comparatively understudied, although recent morphological, transcriptomic, epigenomic, and functional studies have begun to identify candidate regulatory genes. *Cannabis* produces bulbous, sessile, and stalked glandular trichomes, with stalked glandular trichomes on pistillate bracts and calyces representing the dominant cannabinoid-rich structures. These stalked trichomes are morphologically and metabolically distinct from sessile trichomes, with larger secretory heads, 12–16 secretory disc cells, blue-shifted autofluorescence, and cannabinoid– and monoterpene-rich metabolite profiles (Livingston et al., 2020; Tanney et al., 2021; Hancock et al., 2024). Several transcription factors have now been implicated in cannabis glandular trichome development or trichome-associated metabolism. *CsMIXTA*, an R2R3-MYB transcription factor, is highly expressed in female flowers and isolated trichomes, is positively correlated with cannabinoid biosynthetic gene expression during flower development, and increases glandular trichome density, size, and branching when overexpressed in tobacco (Haiden et al., 2022). A MeJA-responsive regulatory pathway has also been proposed, with *CsMYC4* identified as a positive regulator of glandular trichome formation and associated cannabinoid accumulation (Huang et al., 2024). Epigenomic profiling has further shown that cannabis glandular trichomes contain active chromatin marks at cannabinoid and terpenoid biosynthetic genes and metabolite transporters, while trichome-specific H3K56ac-enriched cis-regulatory regions are enriched for transcription factor binding motifs, including MYB-related motifs linked to the strongly trichome-enriched candidate *CsEOBI* (Conneely et al., 2024). More recently, a *CsYABBY3-CsAS1* transcriptional module was described in which a FIL/YAB3-family transcription factor and an R2R3-MYB factor form a positive feedback loop that coordinately promotes glandular trichome differentiation and cannabinoid biosynthesis, illustrating the close integration of developmental and metabolic programs in this system (Zhu et al., 2026). Cross-species transcriptome and gene regulatory network analyses in cannabis, hop, and tomato have also begun to distinguish conserved glandular trichome processes from species-specific regulatory programs (Tamiru-Oli et al., 2026).

Despite these advances, existing studies have largely focused on female tissues, isolated trichomes, organ-specific comparisons, developmental time courses, or hormone perturbation responses. No study has yet systematically evaluated candidate trichome development regulators across *C. sativa* floral phenotypes while experimentally decoupling sex chromosome complement from floral sexual identity. Here, we leveraged a previously published sex-reversal RNA-seq dataset (Monthony et al., 2026), focusing on 46 floral transcriptomes from three commercial high-THC genotypes. This subset includes untreated female flowers (FFs, XX), untreated male flowers (MFs, XY), induced male flowers on XX plants (IMFs), and induced female flowers on XY plants (IFFs), allowing us to distinguish expression patterns associated with chromosomal sex from those associated with the phenotypic and morphological identity of the flower. We hypothesize that cannabis trichome development involves a conserved set of regulatory genes, but that their expression is modulated by phenotypic floral sex. To test this, we compiled an orthology and literature-informed reference set of trichome development-related genes, including regulators characterized in *A. thaliana* and *S. lycopersicum* as well as recently reported candidate trichome regulators from *C. sativa*. We then identified and curated cannabis trichome development-related genes (CsTDRGs), built a stage-specific regulatory model, and compared CsTDRG expression across untreated and sex-reversed floral phenotypes.

## Materials & Methods

### Orthology analysis and pathway modelling

A list of canonical TDRGs was compiled from *A. thaliana* and *S. lycopersicum*. Proteomes used for orthology analysis with OrthoFinder (v2.5.5), were obtained from the NCBI (*A. thaliana:* TAIR10.1; *C. sativa* ‘Pink Pepper’: ASM2916894v1), the Sol Genomics Network (*S. lycopersicum*: ITAG4.0), and from Carey et al. (2026; C. sativa “Otto II”) and analyzed following the approach described in Monthony et al. (2024). Orthologs were identified across two *C. sativa* genome assemblies: the current XX reference genome (‘Pink Pepper’; RefSeq GCF_029168945.1 and a fully phased diploid assembly ‘Otto II’ (XX + XY; Carey et al., 2024), allowing for assignment of CsTDRGs to X-linked, Y-linked, or autosomal loci. Genomic positions of all CsTDRGs were visualized across pseudo-chromosomes using ChromoMap (v4.1.1; Anand & Rodriguez Lopez, 2022). To place the identified CsTDRGs orthologs into a developmental framework, a regulatory pathway model was constructed by integrating functional annotations from the published trichome literature with the orthology results.

### Plant material and RNA-seq data acquisition

Gene expression data were obtained from immature floral tissues sampled 14 days after the onset of floral induction. The dataset, fully described in Monthony *et al*. (2026), includes three high-THC (“drugtype”) *C. sativa* varieties: Deadly Kernel (DK), La Rosca (LR), and Panama Pupil (PP). All experimental plants were propagated as clonal cuttings from verified XX or XY parent plants of each genotype, thereby ensuring identical genetic backgrounds across treatments and replicates.

Four distinct floral phenotypes were generated: (1) Untreated female flowers (FF, chromosomal sex XX, phenotypic sex female). (2) Untreated male flowers (MF, chromosomal sex XY, phenotypic sex male). (3) Induced male flowers on genetically female plants (IMF; chromosomal sex XX, phenotypic sex male) produced by foliar application of silver thiosulfate (STS), which suppresses ethylene signaling and promotes the formation of staminate flowers. (4) Induced female flowers on genetically male plants (IFF; chromosomal sex XY, phenotypic sex female) produced by foliar application of ethephon, which elevates endogenous ethylene levels and promotes the formation of pistillate flowers.

Total RNA was extracted from immature (14-day-old) flowers and RNA-seq libraries were sequenced on an AVITI platform (Element Biosciences, San Diego, California) using 2 × 150 bp paired-end chemistry at the Genomic Analysis Platform of the Institut de biologie intégrative et des systèmes (IBIS), Université Laval, QC, Canada. Reads were quality-trimmed with Trimmomatic, screened for contaminants, aligned to the *C. sativa* ‘Pink Pepper’ v1 and ‘Otto II’ genomes with STAR (v2.7.11b), and gene counts were obtained using HTSeq (v2.0.2).

### Differential expression analysis of CsTDRGs

Differential gene expression (DGE) analysis was performed in R (v4.3.3) using DESeq2 (v1.40.2). Low-count genes were filtered, and variance-stabilizing transformation (VST) was applied for exploratory visualization. Five biologically informative contrasts were evaluated (Figure 2). Three primary pairwise contrasts were used to identify CsTDRGs associated with floral phenotype across untreated and sex-reversed flowers: (C1) untreated male flowers (MF, XY) versus untreated female flowers (FF, XX), representing baseline sexual dimorphism; (C2) induced male flowers (IMF, XX) versus untreated female flowers (FF, XX), representing phenotypic masculinization within the XX background; and (C3) untreated male flowers (MF, XY) versus induced female flowers (IFF, XY), representing phenotypic feminization within the XY background. As a supplementary concordance analysis, (C4) IMF versus IFF was used to test whether candidate CsTDRGs identified in the primary contrast framework retained the expected direction of expression between the two induced floral phenotypes. Finally, (C5) an aggregate floral phenotype contrast compared staminate flowers (MF + IMF) with pistillate flowers (FF + IFF), irrespective of chromosomal background (Figure 2). Log_2_ fold-changes (LFC) were shrunk using *ashr* to reduce noise in effect size estimation. Genes were considered differentially expressed at padj ≤ 0.05 and |log_2_FC| ≥ 1 (Monthony et al., 2026). To ensure robustness and biological reproducibility, only genes consistently differentially expressed across all three genotypes were retained for downstream analyses. All plots were generated using *ggplot2*.

**Figure 2.**
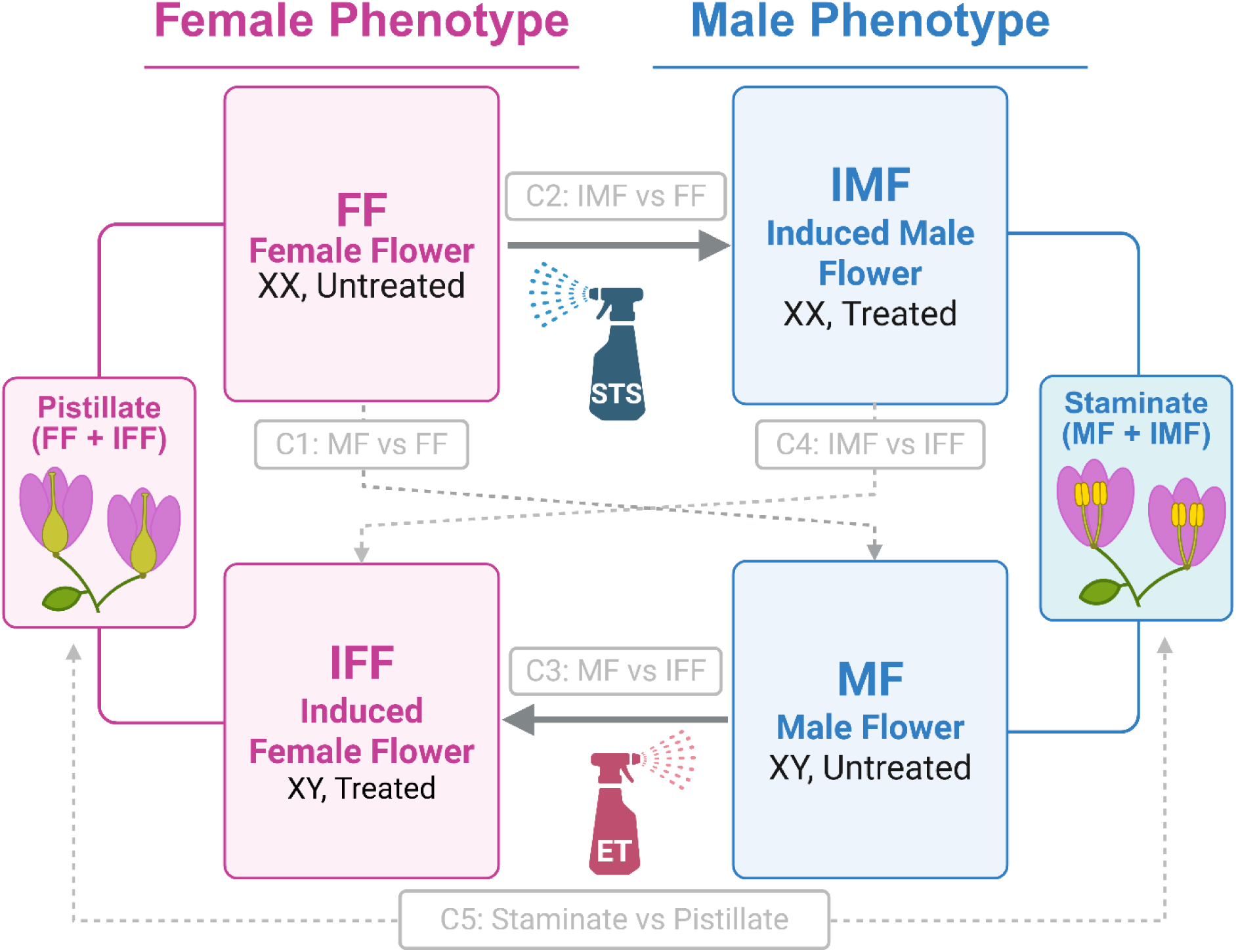
Experimental design used to decouple chromosomal sex from floral phenotype in *C. sativa*. Untreated XX plants produced female flowers (FF), whereas untreated XY plants produced male flowers (MF). Silver thiosulfate (STS) treatment induced male flowers on XX plants (IMF), while ethephon treatment induced female flowers on XY plants (IFF). This design enabled pair-wise comparisons of baseline sexual dimorphism, masculinization within the XX background, feminization within the XY background, and aggregate floral phenotype effects by comparing staminate flowers (MF + IMF) with pistillate flowers (FF + IFF). Grey solid arrows indicate chemical sex-reversal treatments and differential-expression contrasts; dashed arrows indicate differential-expression contrasts only.

## Results

To assess whether candidate trichome development regulator genes (CsTDRGs) are associated with chromosomal sex or floral phenotype, we analyzed their expression across four floral classes: untreated female flowers from XX plants (FF), untreated male flowers from XY plants (MF), induced male flowers from XX plants (IMF), and induced female flowers from XY plants (IFF). This framework allowed CsTDRG expression patterns to be compared between genetic sex and phenotypic floral sex across cultivars.

### Identification and genomic distribution of candidate trichome developmental regulators in *C. sativa*

Orthology-based screening identified 53 unique *C. sativa* candidate trichome development regulator genes (CsTDRGs; Supplementary Table 1). These genes were predominantly autosomal and unevenly distributed across the genome, with the highest numbers on chromosomes 1 and 5 (Figure 3A). Only three CsTDRGs were located on the sex chromosomes; in each case, X– and Y-linked homologs were present in the recombining region (*CsCYCB2, CsTT8* and *CsKAK*).

### Sexual phenotype–associated structure in TDRG expression

Principal component analysis (PCA) of CsTDRG expression at 14 days post-flower induction showed partial separation among floral phenotypes (Figure 3B). PC1 explained 60.85% of the variance and separated IMF and MF samples, whereas FF and IFF samples occupied similar regions of the ordination. PC2 explained 10.23% of the variance and contributed additional separation among samples. Overall, FF and IFF showed greater overlap than the two phenotypically male classes.

**Figure 3.**
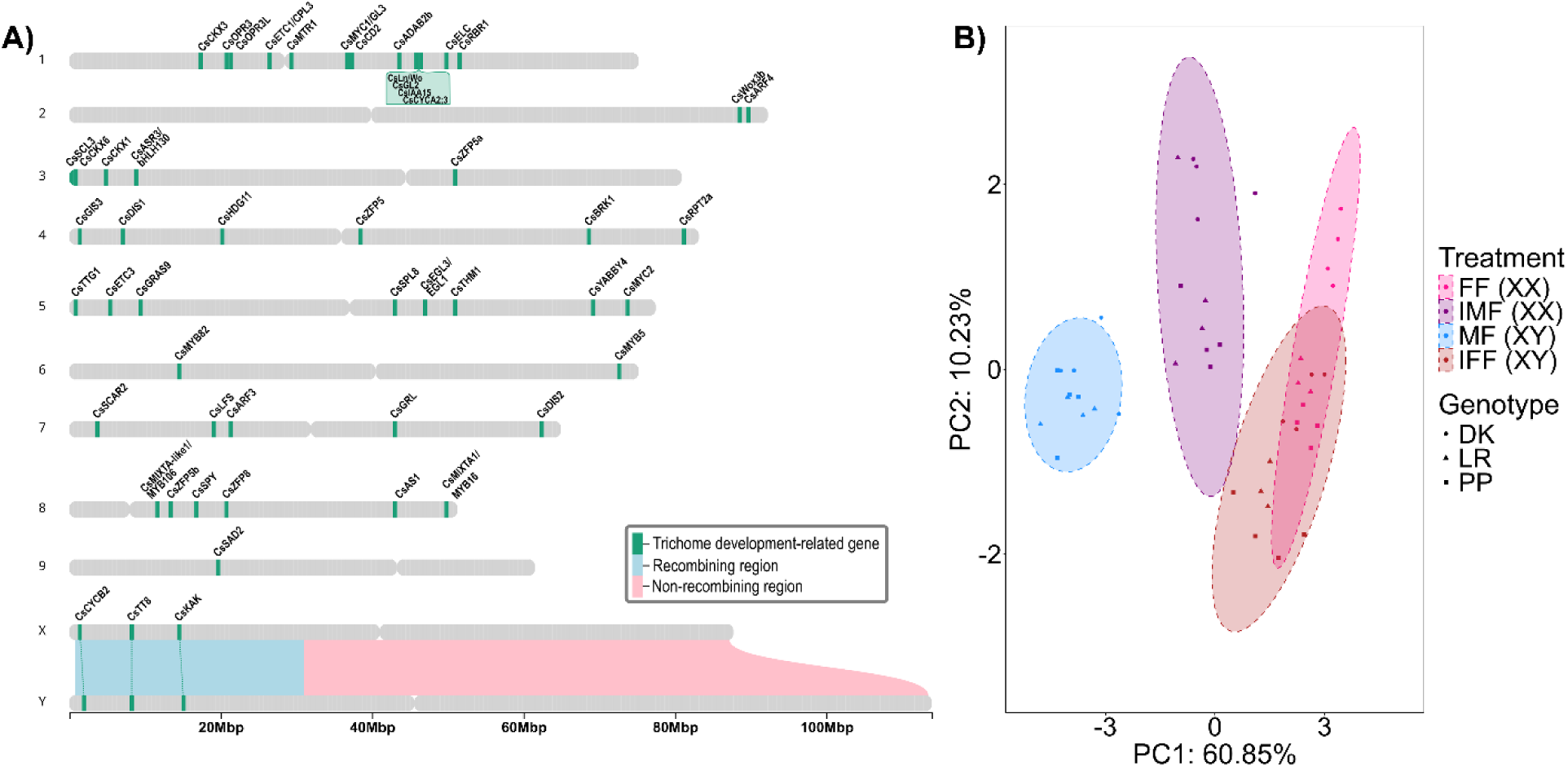
Genomic distribution and expression structure of candidate CsTDRGs. A) Chromosomal localization of CsTDRGs in *C. sativa*. Autosomal loci are shown using the XX reference genome *C. sativa* ‘Pink Pepper’. Recombining and non-recombining regions of the sex chromosomes are shown based on Carey et al. (2024) in *C. sativa* ‘Otto II’. B) PCA of normalized CsTDRG expression across four floral classes at 14 days post-flower induction. FF = female flowers (XX), IMF = induced male flowers (XX), MF = male flowers (XY), and IFF = induced female flowers (XY). Points represent individual samples; shapes denote cultivar.

### Pairwise differential expression identifies phenotype-associated CsTDRGs

Differential expression analysis of the 52 conserved CsTDRGs across the four floral phenotypes identified 16 differentially expressed genes (DEGs). Pairwise differential-expression analysis across the three primary contrasts identified 16 CsTDRGs that met the significance and effect-size criteria in at least one contrast (Figure 4A, B and Supplementary Table 2). These contrasts included untreated MF vs FF, MF vs IFF within the XY background, and IMF vs FF within the XX back-ground, allowing candidate genes to be evaluated for recurrent directionality across untreated and sex-reversed floral phenotypes.

**Figure 4.**
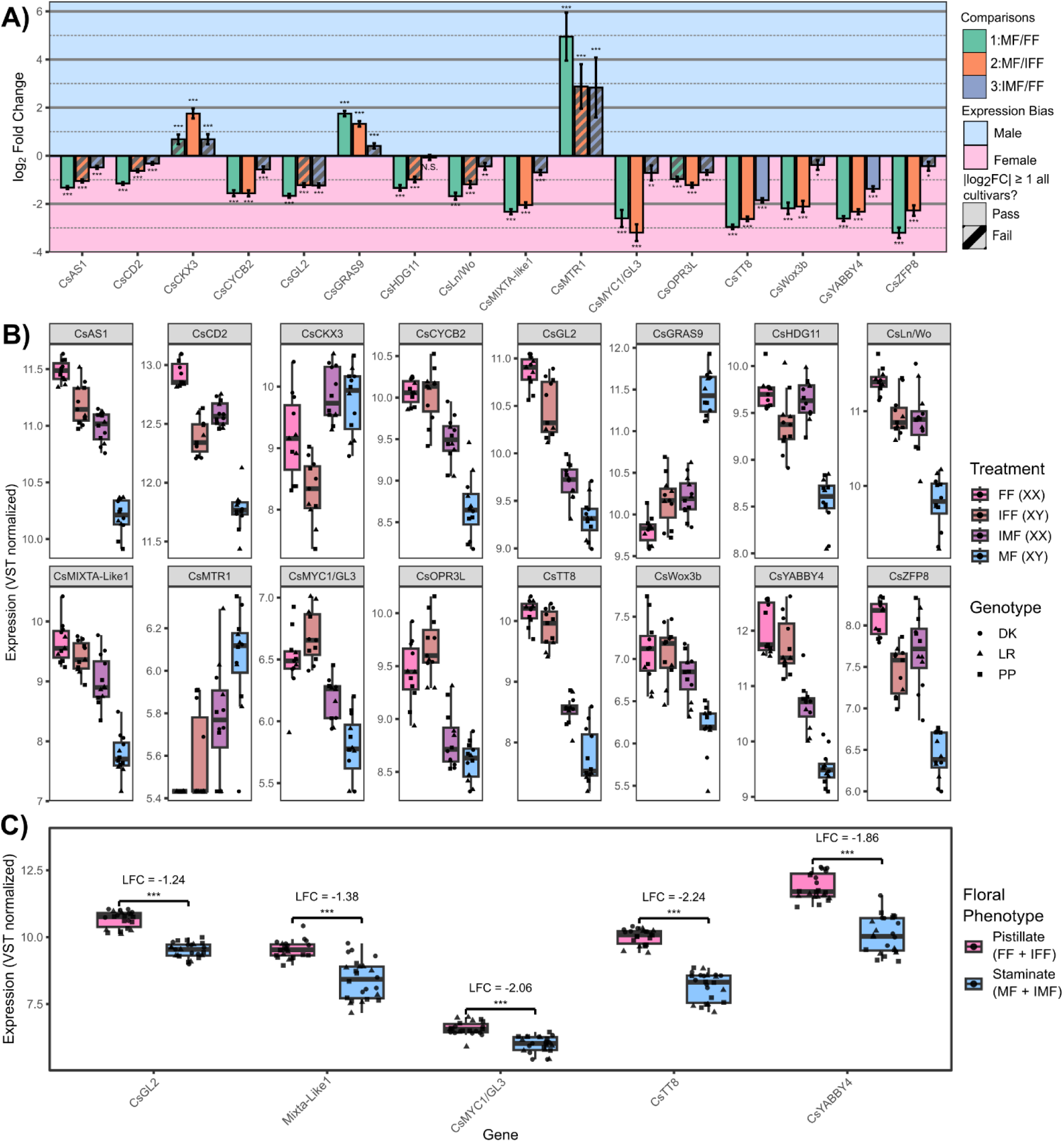
Differential expression of CsTDRGs across floral phenotypes. A) Mean DESeq2 log₂ fold change ± SE for candidate CsTDRGs across three contrasts: MF/FF, MF/IFF, and IMF/FF. Positive values indicate male– or staminate-biased expression; negative values indicate female– or pistillate-biased expression. Stars indicate adjusted p-value thresholds (* padj ≤ 0.05; ** padj ≤ 0.01; *** padj ≤ 0.001; N.S., not significant). Hatched bars indicate genes that did not meet the all-cultivar filter of |log₂FC| ≥ 1 and padj ≤ 0.05. B) VST-normalized expression of candidate CsTDRGs across FF, IFF, IMF, and MF samples. C) VST-normalized expression of CsTDRGs differentially expressed in the pooled pistillate-versus-staminate contrast. Boxplots show pistillate (FF + IFF) and staminate (MF + IMF) samples; points represent biological replicates, with shapes denoting cultivar. Log₂ fold change values and significance levels correspond to the pooled DESeq2 contrast.

In the untreated MF vs FF contrast, 14 CsTDRGs met the significance and effect-size thresholds across all three cultivars, with most showing higher expression in FF. The strongest FF-biased genes included *CsTT8*, *CsYABBY4*, *CsMIXTA-like1/MYB106, CsMYC1/GL3, CsWOX3b*, and *CsZFP8*, while *CsMTR1, CsGRAS9*, and *CsCKX3* represented the main MF-biased candidates (Figure 4A, B; Supplementary Table 2).

The sex-reversal contrasts were then used to assess whether these expression patterns were retained when karyotype and floral phenotype were experimentally decoupled. Several genes showed recurrent female-biased expression across the primary contrast framework, although the strength and cultivar-level consistency varied among genes. *CsTT8* and *CsYABBY4* showed the most consistent female-biased expression, passing the strict all-cultivar threshold in all three contrasts. *CsGL2* was also consistently female-biased across contrasts, although individual cultivar-level effects varied around the log₂FC threshold. Additional genes, including *CsMIXTA-like1/MYB106*, *CsMYC1/GL3*, *CsZFP8*, *CsWOX3b*, *CsLn/Wo*, *CsCYCB2*, and *CsHDG11*, showed strong female-biased expression in MF vs FF and MF vs IFF, but weaker or more cultivar-dependent effects in IMF vs FF. Conversely, *CsMTR1* and *CsGRAS9* were the strongest recurrent male-biased candidates, while *CsCKX3* showed its strongest male-biased expression in the MF vs IFF contrast.

Expression-level visualization across the four floral classes showed corresponding differences in VST-normalized expression for the candidate CsTDRGs (Figure 4B). *CsTT8* and *CsYABBY4* showed higher expression in FF and IFF than in MF and IMF, alongside *CsGL2* and *CsMIXTA-like1/MYB106*, which showed similar female-biased expression patterns. Other candidates, including *CsZFP8*, *CsLn/Wo*, and *CsMYC1/GL3*, showed more intermediate expression in one or both sex-reversed classes. Because the tomato *Lanata* (*SlLn*; Solyc03g031760) and *Woolly* (*SlWo*; Solyc02g080260) queries both resolved to cannabis protein models derived from the same NCBI locus/CDS (XP_030492336.2/XP_030492337.2), this candidate was retained as a single combined entry, *CsLn/Wo*, in downstream analyses.

### Phenotype-level analyses refine candidate CsTDRGs

As a supplementary concordance analysis, the 16 candidate CsTDRGs identified from the primary contrast framework were compared directly between the two induced floral phenotypes (IMF vs IFF). Thirteen of 16 candidates retained the expected direction of effect relative to their average direction across the primary contrasts (Supplementary Table 4). Pistillate-associated candidates including *CsTT8*, *CsGL2*, *CsMIXTA-like1/MYB106*, *CsMYC1/GL3*, and *CsYABBY4* showed higher expression in IFF, whereas *CsCKX3*, *CsGRAS9*, and *CsMTR1* showed higher expression in IMF. Only *CsTT8* and *CsCKX3* passed the strict all-cultivar threshold in this contrast, indicating that IMF vs IFF supports directional concordance rather than serving as an additional discovery screen.

To further summarize CsTDRG expression by floral phenotype, transcriptomes were grouped as pistillate flowers (FF + IFF) or staminate flowers (MF + IMF). Six CsTDRGs met the differential-expression threshold in this pooled contrast (Figure 4C; Supplementary Table 3). Five genes were pistillate-biased: *CsTT8* (log₂FC = −2.24), *CsMYC1/GL3* (log₂FC = −2.06), *CsYABBY4* (log₂FC = −1.86), *CsMIXTA-like1/MYB106* (log₂FC = −1.38), and *CsGL2* (log₂FC = −1.24).

Together, the pairwise, supplementary concordance, and aggregate phenotype analyses identified a focused set of pistillate-biased CsTDRGs, including *CsTT8*, *CsYABBY4*, *CsMYC1/GL3*, *CsGL2*, and *CsMIXTA-like1/MYB106*. Staminate-biased candidates were less consistently recovered across analytical layers, with *CsMTR1*, *CsGRAS9*, and *CsCKX3* identified in the primary contrast framework. Complete contrast-level results are provided in Supplementary Tables 2–4.

## Discussion

### Phenotype-associated trichome regulatory programs in *C. sativa* flowers

This study used chemically induced sexual plasticity to evaluate whether candidate CsTDRGs are associated with chromosomal sex or floral phenotype in *C. sativa*. Across 14-day old floral transcriptomes, pistillate-biased candidates were recovered more consistently than staminate-biased candidates. The focused pistillate-associated set included *CsTT8*, *CsYABBY4*, *CsMYC1/GL3*, *CsGL2*, and *CsMIXTA-like1/MYB106*, whereas staminate-biased candidates were more context-dependent, only *CsMTR1*, *CsGRAS9*, and *CsCKX3* were identified in the treatment level contrast framework. Together, these patterns suggest that, during early trichome development, pistillate flowers preferentially activate a conserved epidermal and trichome regulatory program, whereas staminate flowers express a smaller set of candidate repressors or modulators of trichome developmental competence.

This phenotype-associated expression pattern is biologically relevant because glandular trichomes, particularly capitate-stalked trichomes, are the primary sites of cannabinoid and terpenoid accumulation in *C. sativa* flowers (Livingston et al., 2020; Punja et al., 2023). Female inflorescences develop substantially greater coverage of mature glandular trichomes than male flowers (Huang et al., 2024), yet the regulatory basis of this sexual dimorphism remains underexplored. By focusing on early floral transcriptomes collected during chemically induced sex reversal, our study links this well-known morphological dimorphism to candidate regulatory genes associated with trichome initiation, morphogenesis, and maturation, while separating floral phenotype from sexual karyotype in the experimental design.

Interpreting these candidates requires integrating two partially overlapping trichome-development frameworks, exemplified by *A. thaliana* and *S. lycopersicum*. It remains unclear to what extent the regulatory logic described for unicellular non-glandular trichomes in *A. thaliana* is shared with the more complex multicellular glandular trichomes found in genera such as *Solanum* and *Artemisia* (Chalvin et al., 2020). Because of their simpler structure, unicellular trichomes are often understood through genes with broader roles in epidermal cell fate (Marks, 1997). By contrast, multicellular and glandular trichomes likely rely on these conserved epidermal fate pathways alongside additional mechanisms controlling trichome initiation, morphogenesis, and glandular differentiation (Chalvin et al., 2020; Huchelmann et al., 2017). As such we anticipated shared orthologs between the two model species chosen in our study. Consistent with this expectation, our screening recovered 52 unique CsTDRGs in total, after integrating candidates derived from *Arabidopsis*– and tomato-associated trichome gene sets together with the small number of previously reported cannabis trichome-development genes (Supplementary Table 1). This candidate set therefore provides a useful comparative framework for interpreting cannabis trichome development, but orthology alone cannot resolve the developmental roles of these genes. Functional interpretation requires integrating these orthology relationships with expression patterns across floral phenotype and sexual karyotype.

The genomic distribution of CsTDRGs further supports a regulatory rather than simple structural-genetic explanation for sex-associated trichome dimorphism. Most CsTDRGs were autosomal, and only three candidate genes were located on the sex chromosomes. In each case, X– and Y-linked homologs were present in the recombining region: *CsCYCB2*, *CsTT8*, and *CsKAK* (Figure 3A). Thus, the major expression differences observed between pistillate, and staminate flowers are unlikely to reflect simple presence or absence of candidate trichome genes in the non-recombining sex-determining region. This interpretation is consistent with organ-level observations in monoecious *C. sativa*, where phytocannabinoid-synthesizing glandular trichomes occur on both male and female flowers, but at significantly higher densities in female flowers (Ghosh et al., 2023). Instead, these patterns point toward differential regulation of largely shared developmental genes across floral phenotypes.

### Pistillate-biased CsTDRGs define the strongest phenotype-associated signal

The clearest expression signal was the recurrent enrichment of pistillate-biased CsTDRGs. In the aggregate pistillate-versus-staminate contrast, biological replicates were contrasted by floral phenotype irrespective of chromosomal background (Figure 4C and Supplementary Table 3). Five genes were significantly pistillate-biased: *CsTT8* (log₂FC = −2.24), *CsMYC1/GL3* (log₂FC = −2.06), *CsYABBY4* (log₂FC = −1.86), *CsMIXTA-like1/MYB106* (log₂FC = −1.38), and *CsGL2* (log₂FC = −1.24). Broadly, this pattern is consistent with the greater abundance and density of glandular trichomes reported in pistillate *C. sativa* flowers (Ghosh et al., 2023; Punja et al., 2023) and the reported role of these genes in epidermal fate specification, bHLH/MYB regulatory activity, floral organ identity, and epidermal outgrowth or glandular morphogenesis.

Among these candidates, *CsTT8*, *CsMYC1/GL3*, and *CsGL2* provide the clearest connection to the canonical epidermal fate framework. In *A. thaliana*, trichome cell fate is initiated by TTG1-dependent MBW-related activator complexes comprising an R2R3-MYB transcription factor, a bHLH transcription factor, and the WD40-repeat protein *TTG1*; these complexes transcriptionally activate downstream regulators including *GL2*, which is required for trichome differentiation and maintenance of cell fate (Khosla et al., 2014). *GL3*, *EGL3*, and *TT8* encode related bHLH transcription factors that can function as bHLH components of MBW complexes. Although these bHLH proteins occupy broadly analogous positions within MBW architecture, their developmental outputs are context dependent: *GL3/EGL3* are most strongly associated with trichome initiation, whereas *TT8* is more prominently linked to flavonoid, anthocyanin, and proanthocyanidin regulation, with a minor role in marginal trichome development (Maes et al., 2008). In the present study, *CsTT8* and *CsGL3* were among the most consistent pistillate-biased genes across the primary contrast framework, and both were retained in the aggregate pistillate-versus-staminate contrast. *CsMYC1/GL3* was also pistillate-biased in the aggregate contrast, linking the cannabis candidate set to bHLH regulators with established roles in glandular trichome development and specialized metabolism. In tomato, *SlMYC1* is required for type VI glandular trichome formation and positively regulates monoterpene biosynthesis in glandular trichomes, providing a direct comparative link between bHLH-mediated trichome development and terpene metabolism (Xu et al., 2018). *CsTT8* is similarly notable because *LOC115703590/XM_030630819*, annotated here as *CsTT8* and as *CsbHLH114* in Bassolino *et al*. (2023), was profiled as a bHLH candidate in the context of cannabis anthocyanin regulation. Together, *CsMYC1/GL3* and *CsTT8* highlight pistil-late-biased bHLH candidates that may connect trichome-associated epidermal differentiation with terpene– and flavonoid-associated specialized metabolism. Whether these roles are conserved, rewired, or combined in cannabis glandular trichomes remains to be tested experimentally.

Beyond the MBW-associated candidates, several additional pistillate-biased CsTDRGs point to broader developmental processes that may support glandular trichome formation in female flowers. *CsYABBY4* (LOC115715807; annotated as *CsYABBY3* in Zhu et al., 2026) was retained in the aggregate pistillate-versus-staminate contrast, making it one of the strongest non-MBW candidates in the pistillate-associated set (Figure 4C). This is notable because Zhu et al. functionally characterized the same *C. sativa* locus as a trichome-enriched *FIL/YAB3*-family regulator that promotes glandular trichome formation and cannabinoid accumulation through a positive-feedback module with *CsAS1*, including activation of cannabinoid biosynthetic genes. *CsMIXTA-like1/MYB106* further supports activation of an epidermal outgrowth program in pistillate flowers. MIXTA-like R2R3-MYBs are recurrently implicated in epidermal outgrowth, cuticle formation, and glandular or multicellular trichome development. In tomato, *SlMIXTA-like* promotes trichome formation and directly activates *SlDAHPS*, linking trichome development with primary metabolic flux into the shikimate and phenylpropanoid pathways (Ying et al., 2020). In *A. annua*, *AaMIXTA1* promotes glandular secretory trichome initiation and cuticle biosynthesis (P. Shi et al., 2018; Yan et al., 2018). In cannabis, *CsMIXTA* (LOC115701410) has been identified as a mature-trichome-associated SG9 candidate (Del Rosario-Makridis, 2023) and its overexpression in tobacco has been demonstrated to increase glandular trichome density, size, and branching (Haiden et al., 2022). By contrast, *CsMIXTA-like1* (LOC115698708) has showed lower and less consistent mature-trichome enrichment in prior MYB-focused work (Del Rosario-Makridis, 2023), but was the differentially expressed MIXTA-like paralog retained in the present aggregate pistillate-versus-staminate contrast. This suggests that *CsMIXTA-like1* may be associated with an early pistillate floral or trichome outgrowth context, rather than serving as a general marker of mature glandular trichomes as has been suggested for *CsMIXTA*. Although not retained in the aggregate phenotype contrast, *CsLn/Wo* showed a recurrent pistillate-biased direction across the primary contrast framework and is biologically relevant because tomato *Ln* and *Wo* encode interacting HD-ZIP IV regulators that promote multicellular trichome formation. Although represented here as a combined *CsLn/Wo* orthology assignment because tomato *Ln* and *Wo* resolved to the same cannabis locus in our analysis (Supplementary Table 1), this candidate is notable given that tomato *Ln* and *Wo* cooperatively activate downstream trichome-associated genes, including *SlCycB2* and *SlCycB3* (Xie et al., 2022). The co-recovery of *CsLn/Wo* and *CsCYCB2* in the primary contrast framework is therefore consistent with a possible HD-ZIP IV–CycB-like module contributing to pistillate trichome outgrowth competence (Figure 4 and Supplementary Table *2*).

Additional candidates such as *CsWOX3b* and *CsZFP8* were less consistently retained across analytical layers. *CsWOX3b* demonstrated a strong female-biased pattern of expression in contrasts 1 and 2, which is notable because tomato *SlWox3b* acts downstream of *SlWo* in a dosage-dependent module controlling multicellular trichome type specification (Figure 4A and B). Alongside the female-biased direction of *CsLn/Wo* and the male-biased expression of *CsMTR1*, this pattern aligns with the tomato model in which HD-ZIP IV activity, MTR-mediated feedback, and *WOX3B*-associated programs shape multicellular trichome fate (Wu et al., 2023), identifying this regulatory axis as a priority for future functional study in cannabis. *CsZFP8* may reflect an additional hormone-responsive C2H2 zinc-finger layer of trichome regulation. This candidate is annotated as *ZFP8* in the cannabis reference genome, and during the OrthoFinder analysis grouped with Arabidopsis *GIS/ZFP8-family* regulators as well as tomato *Hair/Hair-like* trichome regulators (Supplementary Table 1). This orthology context is notable because C2H2 zinc-finger regulators have been implicated in trichome development across both unicellular and multicellular systems. In *Arabidopsis*, *GIS/ZFP*-family factors integrate gibberellin and cytokinin signalling upstream of trichome initiation regulators (Sun et al., 2015), while in tomato, *Hair* and *Hair-like* regulate multicellular trichome formation in a jasmonate-responsive pathway involving repression of *THM1* (Hua et al., 2022). In a related tomato C2H2 module, *Hair* also physically interacts with *SlZFP8-like*, and *SlZFP8-like* interacts with *Woolly*, linking *ZFP8*-like activity to multicellular trichome initiation and elongation (Zheng et al., 2022). As broader comparative support, *Artemisia annua AaZFP8L* overexpression increased glandular trichome density and length, elevated terpenoid accumulation, and was linked to ABA biosynthesis, further supporting a connection between *ZFP8*-like regulators, glandular trichome development, phytohormone signalling, and specialized metabolism (Zhang et al., 2024).

### Staminate-biased CsTDRGs implicate repressors and modulators of trichome competence

Given the low glandular trichome abundance of staminate flowers, fewer staminate-biased candidates were expected than pistillate-biased candidates, and this was largely borne out by the data. Staminate-biased candidates were less consistently recovered across analytical layers, with *CsMTR1*, *CsGRAS9*, and *CsCKX3* identified in the treatment-level contrast framework (Figure 2A, B; Supplementary Tables 2). Although these genes operate at different levels of the regulatory hierarchy, they share a broad functional theme across species: each is associated with pathways that can constrain trichome initiation, modify epidermal developmental competence, or alter the hormonal environment in which trichome development occurs. Their enrichment in staminate or phenotypically male floral tissues therefore supports the interpretation that reduced trichome abundance in staminate flowers may reflect not only weaker activation of pistillate-associated trichome-promoting programs, but also preferential expression of putative repressors or modulators of trichome development.

Amongst staminate-associated genes, *CsMTR1* provides the clearest connection to a multicellular trichome regulatory module. In tomato, tomato <u>m</u>ulticellular trichome repressor 1 (*MTR1*) and related MTR proteins act as negative regulators of the HD-ZIP IV transcription factor *Wo*, with loss of *MTR1/MTR2* increasing Wo protein accumulation and shifting trichome development toward long digitate trichomes at the expense of peltate trichomes (Wu et al., 2023, 2024). Thus, the tomato model supports a role for *MTR*-mediated feedback in tuning *Wo* dosage and multicellular trichome fate, rather than a direct positive association between *MTR* expression and glandular trichome identity. In the present study, *CsMTR1* was strongly male-biased across the primary contrast framework (LFC 2.84-4.95; Supplementary Table 1), whereas *CsLn/Wo* and *CsWOX3b* showed female-biased directionality (Figure 2A,B). This reciprocal pattern suggests that cannabis floral phenotypes may differentially regulate an HD-ZIP IV–MTR–WOX axis resembling the module described in tomato multicellular trichomes, with elevated *CsMTR1* in staminate flowers potentially constraining *Wo*-like activity or altering trichome developmental competence rather than directly promoting the glandular trichome program associated with pistillate flowers.

*CsGRAS9* represents a second candidate staminate-biased regulatory component with the potential to constrain trichome-promoting bHLH programs. In tomato, *SlGRAS9* acts as a negative regulator of glandular trichome development by directly binding the *SlMYC1* promoter and repressing its transcription; loss of *SlGRAS9* increases glandular trichome density and elevates *SlMYC1* expression, supporting a repressive *SlGRAS9*–*SlMYC1* relationship (Y. Shi et al., 2025). *SlMYC1* itself is required for type VI glandular trichome formation and positively regulates monoterpene biosynthesis in tomato glandular cells, linking this bHLH-regulated developmental module to specialized metabolism (Xu et al., 2018). In the present study, *CsGRAS9* was male-biased in the primary contrast framework, whereas several bHLH-associated candidates, including *CsTT8* and *CsMYC1/GL3*, were pistillate-biased (Figure 4). Although *CsMYC1/GL3* was identified through the Arabidopsis *MYC1/GL3*-like framework rather than as the direct tomato *SlMYC1* ortholog, the opposing expression of *CsGRAS9* and pistillate-biased bHLH candidates is consistent with the broader possibility that staminate flowers preferentially express regulators that limit trichome-promoting transcriptional programs.

*CsCKX3* may represent a more indirect, hormone-associated component of the staminate-biased signal. Cytokinin oxidase/dehydrogenase genes encode enzymes that irreversibly degrade cytokinins and thereby regulate local cytokinin availability (Schmülling et al., 2003). Although *CKX3* is not a core trichome identity regulator, recent work in tomato linked *CKX1/CKX3* expression and cytokinin homeostasis to multicellular trichome morphogenesis: *CKX1* and *CKX3* were elevated in *SlBRC2a* mutant trichomes, while trichome-targeted *CKX1* expression reduced basal-cell expansion and endoreduplication (Wu et al., 2024). In the present study, *CsCKX3* showed male-biased expression in the primary contrast framework (Figure 4A, B), suggesting that, if a similar mechanism operates in cannabis, cytokinin turnover may contribute to the hormonal environment associated with staminate floral tissues rather than acting as a direct determinant of glandular trichome identity.

Collectively, the staminate-biased candidates suggest that the comparatively low trichome abundance of male flowers may arise not only from reduced activation of pistillate-biased trichome-promoting pathways, but also from preferential expression of regulatory components associated with constrained trichome developmental competence. *CsMTR1*, *CsGRAS9* and *CsCKX3* point to distinct but complementary layers of regulation, including HD-ZIP IV dosage control, repression of bHLH-associated trichome programs and cytokinin homeostasis. Together, these patterns support a model in which sex-associated trichome dimorphism in cannabis reflects differential regulation of shared developmental and hormonal pathways across floral phenotypes.

### A hybrid model for phenotype-associated trichome regulation in cannabis flowers

The expression patterns identified here support a model in which sexually dimorphic trichome development in *C. sativa* is regulated through a shared but differentially activated developmental toolkit. Rather than fitting cleanly into either the canonical *Arabidopsis* model of unicellular non-glandular trichome initiation or the tomato model of multicellular glandular trichome morphogenesis, cannabis appears to combine elements of both. This is reflected in Figure 5, which organizes candidate CsTDRGs into two broad developmental categories: genes associated with trichome fate and outgrowth initiation, and genes associated with subsequent morphogenesis, maturation, and trichome type specification. This framework allows the pistillate– and staminate-biased candidates to be interpreted as components of a broader regulatory hierarchy, in which conserved epidermal fate regulators are integrated with initiation and morphogenesis factors more typically associated with multicellular or glandular trichome systems.

**Figure 5.**
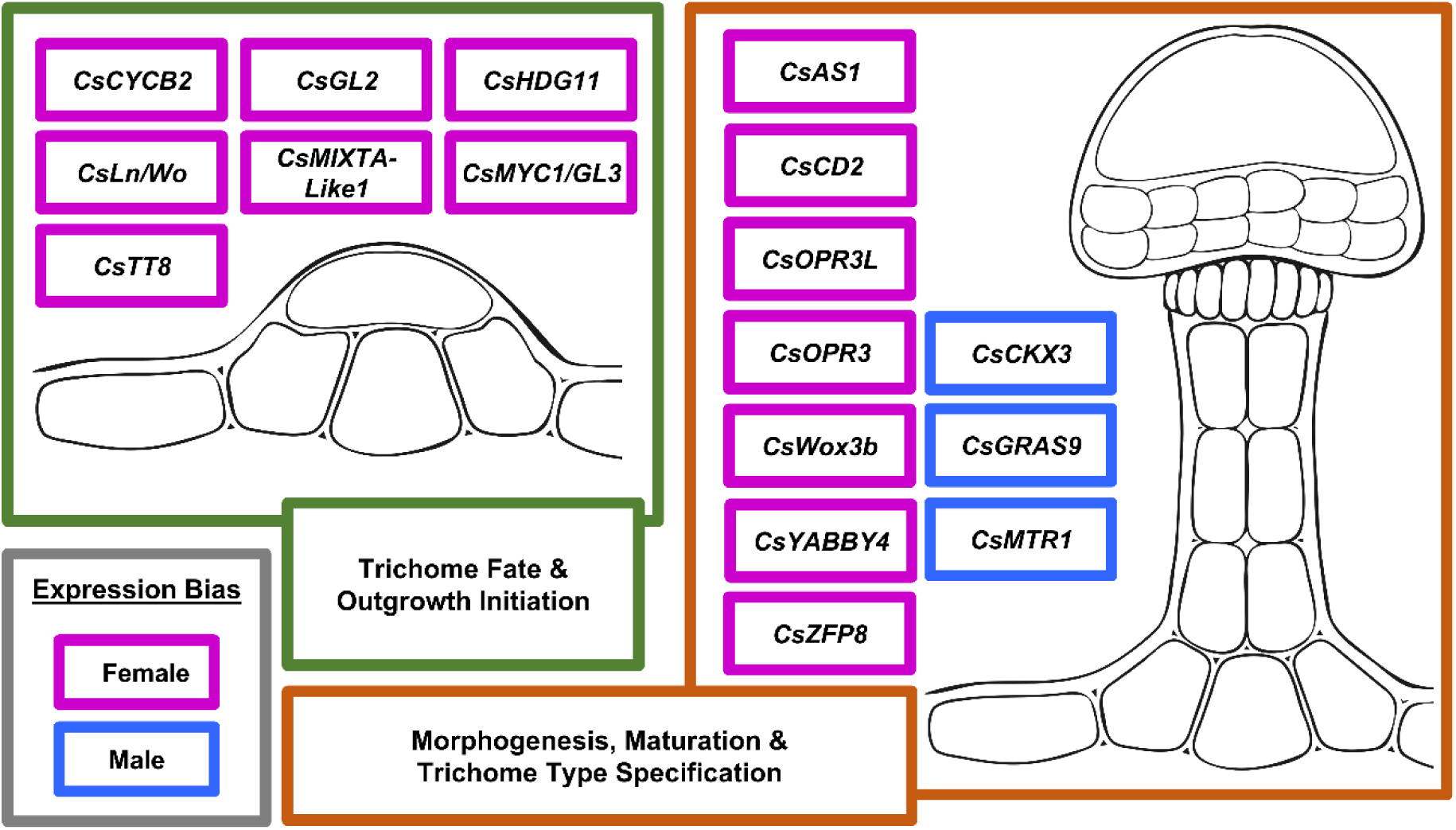
Functional organization of candidate CsTDRGs associated with sexually dimorphic trichome development in *C. sativa*. Candidate CsTDRGs were grouped into two broad developmental categories: trichome fate and outgrowth initiation, and morphogenesis, maturation and trichome type specification. Gene placement reflects inferred primary roles based on comparative evidence from *A. thaliana*, *S. lycopersicum*, *C. sativa*, and other multicellular or glandular trichome systems. Because several regulators may act across multiple stages of trichome development, genes were assigned according to their most well-supported or most frequently reported role. Pink outlines indicate female-biased expression, and blue outlines indicate male-biased expression in the present study.

One indication that cannabis does not simply recapitulate the canonical *Arabidopsis* initiation model is the incomplete modeling of the *GL1/MYB23*-centred MBW framework. In *A. thaliana*, *GL1* acts as a central R2R3-MYB component of the MBW complex that promotes unicellular non-glandular trichome initiation, with *MYB23* providing partially redundant activity and *GL2* acting downstream in trichome cell fate and differentiation (Khosla et al., 2014). In our orthology analysis, clear cannabis orthologs of *GL1* and *MYB23* were not recovered. This negative result should not be interpreted as definitive evidence that cannabis lacks all GL1/MYB23-like activity; however, it is consistent with a more detailed cannabis MYB-family analysis that similarly reported no clear GL1-like R2R3-MYB and suggested that the closest SG15-like candidate was more WER-like than GL1-like, a distinction that is relevant because *WER* is classically associated with root epidermal patterning rather than aerial trichome initiation (Del Rosario-Makridis, 2023). While a putative GL1-like locus (XM_030629461.1) was reported by Garcia-de Heer *et al*. (2025), its annotation has been deprecated in the current NCBI *C. sativa* reference genome annotation, further highlighting the difficulty in identifying this gene through orthology and computation annotation alone.

Evidence from hop, the closest major comparative system within Cannabaceae, also supports caution in expecting a readily identifiable Arabidopsis-like *GL1* node in glandular trichome development. In *Humulus lupulus*, Patzak et al. (2021) did not center lupulin gland development around a clear *GL1* ortholog. Instead, they emphasized alternative MYB-family candidates, including *HlMYB106/MIXTA*, alongside *HlGLABRA2*, *HlCYCB2-4*, *HlZFP8*, and *HlYABBY1*, several of which were more highly expressed in bitter hop cultivars with higher lupulin gland density. This pattern does not demonstrate that *GL1*-like function is absent from Cannabaceae glandular trichomes, but it does suggest that glandular trichome development in this family may rely on a broader or modified epidermal regulatory toolkit rather than a simple *GL1*-centred Arabidopsis module. At the same time, several MBW-related or downstream epidermal fate components were recovered in the present study and showed pistillate-biased expression, including *CsTT8*, *CsMYC1/GL3*, *CsGL2*, and *CsHDG11*. Thus, rather than indicating a wholesale absence of Ara-bidopsis-like regulation, our results point to a modified epidermal fate module in which some components of the MBW framework are retained while others may be absent, highly diverged, or replaced by lineage-specific regulators.

Developmental timing may help explain why different cannabis trichome studies emphasize different candidate regulators. Garcia-de Heer *et al*. (2025) sampled earlier (7 days post-flowering) during ethephon-induced sex reversal and reported female-biased expression of several candidate genes associated with early floral and trichome developmental programs, whereas the present study analyzed 14-day floral tissues during a later stage of altered-sex floral development. Rather than resolving these studies into a single linear sequence, the differences likely reflect that cannabis trichome development is temporally structured: floral identity and early outgrowth signals may precede later programs associated with glandular differentiation, morphogenesis, and maturation. This interpretation is consistent with the sessile-to-stalked transition model described by Livingston *et al*. (2020), in which stalked glandular trichomes on cannabis flowers develop from sessile-like precursors during floral maturation. In that framework, our 14-day floral transcriptomes may capture a developmental window in which trichome fate/outgrowth initiation and early glandular morphogenesis overlap. The recovery of MBW-associated factors, *MIXTA-like1* and HD-ZIP IV regulators, *YABBY/AS1*-associated genes, and hormone-linked modulators is therefore consistent with a transitional regulatory state rather than a single initiation-only or maturation-only phase.

The multicellular and glandular dimensions of the cannabis candidate set further support this hybrid interpretation. Tomato trichome development provides a useful comparative framework because multicellular trichome initiation, morphogenesis, and type specification are strongly associated with HD-ZIP IV, *MIXTA-like*, *WOX*, *MTR*, and hormone-responsive regulators. Several cannabis candidates identified here map onto this broader tomato-like regulatory space, including *CsLn/Wo*, *CsMTR1*, *CsWOX3b*, *CsMIXTA-like1/MYB106*, *CsGRAS9*, and *CsCKX3*. *Artemisia annua* provides an additional useful comparison because glandular trichome initiation involves cooperation between HD-ZIP IV and MIXTA-like regulators, specifically the *AaHD8*–*AaMIXTA1* complex, which promotes *AaHD1* expression and epidermal development, including glandular trichome initiation and cuticle formation (Yan et al., 2018). This comparison is particularly relevant because cannabis also recovered both HD-ZIP IV– and MIXTA-like candidates, suggesting that glandular trichome initiation may involve regulatory combinations not captured by the Arabidopsis MBW model alone. Finally, the recovery of *CsYABBY4 (LOC115715807;* annotated as *CsYABBY3* in *Zhu et al.*, 2026) highlights a cannabis-specific or lineage-modified component not easily placed within either the Arabidopsis or tomato frameworks. Zhu *et al*. (2026) functionally linked this locus to glandular trichome formation and cannabinoid biosynthesis through a *CsYABBY3-CsAS1* module, suggesting that cannabis may integrate floral organ identity, glandular differentiation, and specialized metabolism within its trichome regulatory program.

The altered-sex design adds an additional layer to this model by separating floral phenotype from sexual karyotype. Most CsTDRGs were autosomal, and the few sex-linked candidates were located in recombining regions rather than being exclusive to the non-recombining sex-determining region. Together with the directional concordance observed in sex-reversed flowers, this supports the interpretation that sexually dimorphic trichome development depends largely on differential regulation of a shared CsTDRG repertoire across floral phenotypes. However, this does not exclude an upstream role for sex-chromosome-linked or sex-determination-associated regulators, as has recently been suggested in cannabis (Carey et al., 2026; Garcia-de Heer et al., 2025). Rather, sex-linked cues may establish floral identity or hormonal states that subsequently regulate autosomal and PAR-associated trichome developmental programs. In this sense, trichome abundance and glandular differentiation may behave as downstream sex-associated traits: not encoded by sex-specific trichome genes themselves but activated or constrained by broader floral-sex regulatory networks.

Several limitations should be acknowledged. First, orthology-based discovery is anchored on *A. thaliana* and *S. lycopersicum* and is unlikely to recover cannabis-specific or rapidly diverging regulators. Second, the analysis is restricted to a single developmental window (14 days postinduction) and to whole floral tissue, limiting resolution across initiation, morphogenesis, and maturation; integration with earlier sampling (Garcia-de Heer et al., 2025) and trichome-enriched datasets (Conneely et al., 2024; Tamiru-Oli et al., 2026) will refine stage-specific assignments. Third, chemical sex reversal with STS and ethephon perturbs hormone signaling and may contribute to phenotype-associated differences, particularly for *CsCKX3* and *CsZFP8*. Spatial or single-cell profiling, time-course sampling across the sessile-to-stalked transition, and CRISPR/Cas9 perturbation of prioritized CsTDRGs will be needed to test the hybrid Arabidopsis–tomato model proposed here.

Overall, the model emerging from this study is that pistillate cannabis flowers preferentially activate trichome-promoting regulatory programs that integrate epidermal fate, multicellular out-growth, glandular differentiation, and specialized metabolism, whereas staminate flowers show weaker activation of these programs and higher expression of selected modulators or repressors of trichome developmental competence. This synthesis prioritizes *CsTT8*, *CsMYC1/GL3*, *CsGL2*, *CsMIXTA-like1/MYB106*, *CsYABBY4*, *CsLn/Wo*, *CsMTR1*, *CsGRAS9*, and *CsCKX3* as candidates for future functional testing. Spatial expression analysis, trichome-stage-specific transcriptomics, and targeted perturbation of these candidates will be needed to determine how floral phenotype regulates the developmental and metabolic programs underlying sexually dimorphic trichome production.

## Acknowledgments

The authors gratefully acknowledge the support of the Natural Sciences and Engineering Research Council (NSERC) Discover Grant number RGPIN-2022-03396. ASM has also been supported by a NSERC Canada Vanier Graduate Scholarship. JR is supported by the FRQ Bourse de maîtrise en recherche 2026-2027, https://doi.org/10.69777/2009140 TJ research internship was generously supported by the Université Catholique de Lyon École d’ingénieurs en biotechnologies as well as by the Bourse région mobilité interne from the Région Auvergne-Rhône-Alpes.

## Author Contributions

Conceptualization: ASM, DT

Data Curation: TJ, ASM

Formal Analysis: TJ, JR, ASM, MN

Funding Acquisition: DT

Investigation: TJ, ASM, JR

Methodology: TJ, ASM, JR

Project Administration: ASM, DT

Resources: DT

Software: JR, TJ, ASM

Supervision: ASM, DT

Visualization: JR, TJ, ASM

Writing-Original Draft Preparation: TJ, ASM, MN

Writing-Review & Editing: ASM, MN, JR, DT

## Data Availability Statement

RNA-seq data analyzed in this study have been deposited in the NCBI Sequence Read Archive (SRA) and are publicly available under BioProject accession PRJNA1404156.

## Supplementary Table Captions

**Supplementary Table 1** C. sativa TDRG (trichome development related gene) as identified through a review of existing C. sativa trichome literature and homology analysis of model trichome development species A. thaliana and S. lycopersicum (Homolog IDs: A. thaliana = AT, or S. lycopersicum = Sl) using the current reference genome (‘Pink Pepper’; NCBI, 2023), X and Y haplotypes from the Otto II genotype (Carey et al., 2024b). Chr: chromosome number. Scaf: gene mapped to unplaced scaffold.

**Supplementary Table 2** Differential expression as calculated across differentially expressed candidate CsTDRGs. L2FC = Log2Fold Change; Avg = mean L2FC across all 3 cultivars tested (DK= Deadly Kernel; LR = La Rosca; PP = Panama Pupil V4); SE = Standard Error of the mean of L2FC; Significance column represents adjusted p-values thresholds: * padj ≤ 0.05; ** padj ≤ 0.01; *** padj ≤ 0.001; N.S., not significant.

**Supplementary Table 3** Differential expression of CsTDRGs in the aggregate staminate-versus-pistillate floral phenotype contrast. Differential expression was calculated by contrasting staminate flowers (MF + IMF) against pistillate flowers (FF + IFF), irrespective of chromosomal background. Genes shown met the differential-expression threshold of |log₂FC| ≥ 1 and adjusted p ≤ 0.05. Log₂ fold-change values are reported as staminate relative to pistillate, such that positive values indicate higher expression in staminate flowers and negative values indicate higher expression in pistillate flowers. Results are shown for the average phenotype-level contrast and for individual cultivars. DK = Deadly Kernel; LR = La Rosca; PP = Panama Pupil V4. Significance codes indicate adjusted p-value thresholds: * adjusted p ≤ 0.05; ** adjusted p ≤ 0.01; *** adjusted p ≤ 0.001.

**Supplementary Table 4** Directional concordance of candidate CsTDRGs in the induced male-versus-induced female contrast. Differential expression was calculated for induced male flowers (IMF, XX) versus induced female flowers (IFF, XY) for the 16 candidate CsTDRGs identified in the primary pairwise contrast framework. Log₂ fold-change values are reported as IMF relative to IFF, such that positive values indicate higher expression in induced male flowers and negative values indicate higher expression in induced female flowers. The expected IMF versus IFF bias was inferred from the average log₂ fold change across the primary contrasts. The table reports average values across the three cultivars used in the study, with significance codes indicating adjusted p-value thresholds: N.S., not significant; * adjusted p ≤ 0.05; ** adjusted p ≤ 0.01; *** adjusted p ≤ 0.001. Concordance indicates whether the observed IMF versus IFF direction matched the expected direction from the primary contrast framework.

## References

1. Anand, L., & Rodriguez Lopez, C. M. (2022). ChromoMap: An R package for interactive visualization of multiomics data and annotation of chromosomes. BMC Bioinformatics, 23(1), 33. 10.1186/s12859-021-04556-z

2. Andre, C. M., Hausman, J. F., & Guerriero, G. (2016). Cannabis sativa: The plant of the thousand and one molecules. Frontiers in Plant Science, 7(FEB2016), 1–17. 10.3389/fpls.2016.00019

3. Bassolino, L., Fulvio, F., Pastore, C., Pasini, F., Gallina Toschi, T., Filippetti, I., & Paris, R. (2023). When Cannabis sativa L. Turns Purple: Biosynthesis and Accumulation of Anthocyanins. Antioxidants, 12(7), 1393. 10.3390/antiox12071393

4. Carey, S. B., Bentz, P. C., Lovell, J. T., Akozbek, L. M., Myers, Z. A., Korani, W., Havill, J. S., Padgitt-Cobb, L., Lynch, R. C., Allsing, N., Mangels, J., Stansell, Z., Stack, G. M., Gordon, T., Osmanski, A., Easterling, K. A., Orozco, L. R., Marcus, Z. E., Hale, H., … Harkess, A. (2026). An X-linked sex determination mechanism in cannabis and hop. Nature Communications. 10.1038/s41467-026-73233-7

5. Chalvin, C., Drevensek, S., Dron, M., Bendahmane, A., & Boualem, A. (2020). Genetic Control of Glandular Trichome Development. Trends in Plant Science, 25(5), 477–487. 10.1016/j.tplants.2019.12.025

6. Chang, J., Yu, T., Yang, Q., Li, C., Xiong, C., Gao, S., Xie, Q., Zheng, F., Li, H., Tian, Z., Yang, C., & Ye, Z. (2018). *Hair*, encoding a single C2H2 zincfinger protein, regulates multicellular trichome formation in tomato. The Plant Journal, 96(1), 90–102. 10.1111/tpj.14018

7. Del Rosario-Makridis, G. N. (2023). Molecular Regulation of Glandular Trichome Initiation and Morphogenesis in Cannabis sativa L.

8. Flajšman, M., Slapnik, M., & Murovec, J. (2021). Production of Feminized Seeds of High CBD Cannabis sativa L. by Manipulation of Sex Expression and Its Application to Breeding. Frontiers in Plant Science, 12(November), 1–12. 10.3389/fpls.2021.718092

9. Garcia-de Heer, L., Guo, Q., Mieog, J., Nolan, M., Liu, L., Dimopoulos, N., Melzer, R., & Kretzschmar, T. (2025). A transcriptomic analysis of ethephon-induced sex reversion of male Cannabis sativa L. reveals changes in expression of floral homeotic genes and a distinct trichome morphology. Journal of Experimental Botany. 10.1093/jxb/eraf291

10. Ghosh, D., Chaudhary, N., Shanker, K., Kumar, B., & Kumar, N. (2023). Monoecious Cannabis sativa L. discloses the organ-specific variation in glandular trichomes, cannabinoids content and antioxidant potential. Journal of Applied Research on Medicinal and Aromatic Plants, 35, 100476. 10.1016/j.jarmap.2023.100476

11. Haiden, S. R., Apicella, P. V., Ma, Y., & Berkowitz, G. A. (2022). Overexpression of CsMIXTA, a Transcription Factor from Cannabis sativa, Increases Glandular Trichome Density in Tobacco Leaves. Plants, 11(11), 1519. 10.3390/plants11111519

12. Hancock, J., Livingston, S. J., & Samuels, L. (2024). Building a biofactory: Constructing glandular trichomes in Cannabis sativa. Current Opinion in Plant Biology, 80, 102549. 10.1016/j.pbi.2024.102549

13. Hanuš, L. O., Meyer, S. M., Muñoz, E., Taglialatela-Scafati, O., & Appendino, G. (2016). Phytocannabinoids: A unified critical inventory. Natural Product Reports, 33(12), 1357–1392. 10.1039/C6NP00074F

14. Hua, B., Chang, J., Han, X., Xu, Z., Hu, S., Li, S., Wang, R., Yang, L., Yang, M., Wu, S., Shen, J., Yu, X., & Wu, S. (2022). H and HL synergistically regulate jasmonatetriggered trichome formation in tomato. Horticulture Research, 9, uhab080. 10.1093/hr/uhab080

15. Hua, B., Chang, J., Wu, M., Xu, Z., Zhang, F., Yang, M., Xu, H., Wang, L., Chen, X., & Wu, S. (2021). Mediation of JA signalling in glandular trichomes by the *woolly/SlMYC1* regulatory module improves pest resistance in tomato. Plant Biotechnology Journal, 19(2), 375–393. 10.1111/pbi.13473

16. Huang, X., Chen, W., Zhao, Y., Chen, J., Ouyang, Y., Li, M., Gu, Y., Wu, Q., Cai, S., Guo, F., Zhu, P., Ao, D., You, S., Vasseur, L., & Liu, Y. (2024). Deep learning-based quantification and transcriptomic profiling reveal a methyl jasmonatemediated glandular trichome for-mation pathway in *Cannabis sativa*. The Plant Journal, 118(4), 1155–1173. 10.1111/tpj.16663

17. Huchelmann, A., Boutry, M., & Hachez, C. (2017). Plant Glandular Trichomes: Natural Cell Factories of High Biotechnological Interest. Plant Physiology, 175(1), 6–22. 10.1104/pp.17.00727

18. Ilgenfritz, H., Bouyer, D., Schnittger, A., Mathur, J., Kirik, V., Schwab, B., Chua, N.-H., Jürgens, G., & Hülskamp, M. (2003). The Arabidopsis *STICHEL* Gene Is a Regulator of Trichome Branch Number and Encodes a Novel Protein. Plant Physiology, 131(2), 643–655. 10.1104/pp.014209

19. Khosla, A., Paper, J. M., Boehler, A. P., Bradley, A. M., Neumann, T. R., & Schrick, K. (2014). HD-Zip Proteins GL2 and HDG11 Have Redundant Functions in Arabidopsis Trichomes, and GL2 Activates a Positive Feedback Loop via MYB23[W]. The Plant Cell, 26(5), 2184–2200. 10.1105/tpc.113.120360

20. Livingston, S. J., Quilichini, T. D., Booth, J. K., Wong, D. C. J., Rensing, K. H., Laflamme-Yonkman, J., Castellarin, S. D., Bohlmann, J., Page, J. E., & Samuels, A. L. (2020). Cannabis glandular trichomes alter morphology and metabolite content during flower maturation. The Plant Journal, 101(1), 37–56. 10.1111/tpj.14516

21. Maes, L., Inzé, D., & Goossens, A. (2008). Functional Specialization of the TRANSPARENT TESTA GLABRA1 Network Allows Differential Hormonal Control of Laminal and Marginal Trichome Initiation in Arabidopsis Rosette Leaves. Plant Physiology, 148(3), 1453–1464. 10.1104/pp.108.125385

22. Marks, M. D. (1997). Molecular Genetic Analysis of Trichome Development in Arabidopsis. Annual Review of Plant Physiology and Plant Molecular Biology, 48(1), 137–163. 10.1146/annurev.arplant.48.1.137

23. Monthony, A. S., de Ronne, M., & Torkamaneh, D. (2024). Exploring ethylene-related genes in Cannabis sativa: Implications for sexual plasticity. Plant Reproduction. 10.1007/s00497-023-00492-5

24. Monthony, A. S., Roy, J., De Ronne, M., Carlson, O., Murch, S. J., & Torkamaneh, D. (2026). Sex-specific ethylene responses drive floral sexual plasticity in *Cannabis sativa*. The Plant Journal, 125(3), e70721. 10.1111/tpj.70721

25. Mudge, E. M., Brown, P. N., & Murch, S. J. (2019). The Terroir of *Cannabis*: Terpene Metabolomics as a Tool to Understand *Cannabis sativa* Selections. Planta Medica, 85(9–10), 781–796. 10.1055/a-0915-2550

26. Mudge, E. M., Murch, S. J., & Brown, P. N. (2018). Chemometric Analysis of Cannabinoids: Chemotaxonomy and Domestication Syndrome. Scientific Reports, 8(1), 1–9. 10.1038/s41598-018-31120-2

27. Patzak, J., Henychová, A., & Matoušek, J. (2021). Developmental regulation of lupulin gland-associated genes in aromatic and bitter hops (Humulus lupulus L.). BMC Plant Biology, 21(1), 534. 10.1186/s12870-021-03292-z

28. Punja, Z. K., Sutton, D. B., & Kim, T. (2023). Glandular trichome development, morphology, and maturation are influenced by plant age and genotype in high THC-containing cannabis (Cannabis sativa L.) inflorescences. Journal of Cannabis Research, 5(1), 12. 10.1186/s42238-023-00178-9

29. Raman, V., Lata, H., Chandra, S., Khan, I. A., & ElSohly, M. A. (2017). Morpho-Anatomy of Marijuana (Cannabis sativa L.). In S. Chandra, H. Lata, & M. A. ElSohly (Eds.), Cannabis sativa L.-Botany and Biotechnology (pp. 123–136). Springer International Publishing. 10.1007/978-3-319-54564-6_5

30. Schmülling, T., Werner, T., Riefler, M., Krupková, E., & Bartrina Y, Manns, I. (2003). Structure and function of cytokinin oxidase/dehydrogenase genes of maize, rice, Arabidopsis and other species. Journal of Plant Research, 116(3), 241–252. 10.1007/s10265-003-0096-4

31. Shi, P., Fu, X., Shen, Q., Liu, M., Pan, Q., Tang, Y., Jiang, W., Lv, Z., Yan, T., Ma, Y., Chen, M., Hao, X., Liu, P., Li, L., Sun, X., & Tang, K. (2018). The roles of *Aa MIXTA 1* in regulating the initiation of glandular trichomes and cuticle biosynthesis in *Artemisia annua*. New Phytologist, 217(1), 261–276. 10.1111/nph.14789

32. Shi, Y., Wang, Y., Pan, Y., Deng, C., Zeng, T., Su, D., Lu, W., Lin, Y., Han, J., Deng, W., Wu, S., Liu, Y., Li, N., Li, J., Dong, B., Abid, G., Bouzayen, M., Pirrello, J., Li, Z., & Huang, B. (2025). The SlGRAS9-SlMYC1 regulatory module controls glandular trichome formation and modulates resilience to pest in tomato. The Plant Journal, 122(3), e70183. 10.1111/tpj.70183

33. Song, S., Liu, B., Song, J., Pang, S., Song, T., Gao, S., Zhang, Y., Huang, H., & Qi, T. (2022). A molecular framework for signaling crosstalk between jasmonate and ethylene in anthocyanin biosynthesis, trichome development, and defenses against insect herbivores in *Arabidopsis*. Journal of Integrative Plant Biology, 64(9), 1770–1788. 10.1111/jipb.13319

34. Sun, L., Zhang, A., Zhou, Z., Zhao, Y., Yan, A., Bao, S., Yu, H., & Gan, Y. (2015). *GLABROUS INFLORESCENCE STEMS* (*GIS3*) regulates trichome initiation and development in *Arabidopsis*. New Phytologist, 206(1), 220–230. 10.1111/nph.13218

35. Tanney, C. A. S., Backer, R., Geitmann, A., & Smith, D. L. (2021). Cannabis Glandular Trichomes: A Cellular Metabolite Factory. Frontiers in Plant Science, 12, 721986. 10.3389/fpls.2021.721986

36. Tetali, S. D. (2019). Terpenes and isoprenoids: A wealth of compounds for global use. Planta, 249(1), 1–8. 10.1007/s00425-018-3056-x

37. Wang, S., Barron, C., Schiefelbein, J., & Chen, J. (2010). Distinct relationships between GLA-BRA2 and single-repeat R3 MYB transcription factors in the regulation of trichome and root hair patterning in Arabidopsis. New Phytologist, 185(2), 387–400. 10.1111/j.1469-8137.2009.03067.x

38. Wu, M., Bian, X., Hu, S., Huang, B., Shen, J., Du, Y., Wang, Y., Xu, M., Xu, H., Yang, M., & Wu, S. (2024). A gradient of the HD-Zip regulator Woolly regulates multicellular trichome morphogenesis in tomato. The Plant Cell, 36(6), 2375–2392. 10.1093/plcell/koae077

39. Wu, M., Chang, J., Han, X., Shen, J., Yang, L., Hu, S., Huang, B.-B., Xu, H., Xu, M., Wu, S., Li, P., Hua, B., Yang, M., Yang, Z., & Wu, S. (2023). A HD-ZIP transcription factor specifies fates of multicellular trichomes via dosage-dependent mechanisms in tomato. Developmental Cell, 58(4), 278–288.e5. 10.1016/j.devcel.2023.01.009

40. Xie, Q., Xiong, C., Yang, Q., Zheng, F., Larkin, R. M., Zhang, J., Wang, T., Zhang, Y., Ouyang, B., Lu, Y., Ye, J., Ye, Z., & Yang, C. (2022). A novel regulatory complex mediated by *Lanata* (*Ln*) controls multicellular trichome formation in tomato. New Phytologist, 236(6), 2294–2310. 10.1111/nph.18492

41. Xu, J., Van Herwijnen, Z. O., Dräger, D. B., Sui, C., Haring, M. A., & Schuurink, R. C. (2018). SlMYC1 Regulates Type VI Glandular Trichome Formation and Terpene Biosynthesis in Tomato Glandular Cells. The Plant Cell, 30(12), 2988–3005. 10.1105/tpc.18.00571

42. Yan, T., Li, L., Xie, L., Chen, M., Shen, Q., Pan, Q., Fu, X., Shi, P., Tang, Y., Huang, H., Huang, Y., Huang, Y., & Tang, K. (2018). A novel HD-ZIP IV/MIXTA complex promotes glandular trichome initiation and cuticle development in *Artemisia annua*. New Phytologist, 218(2), 567–578. 10.1111/nph.15005

43. Ying, S., Su, M., Wu, Y., Zhou, L., Fu, R., Li, Y., Guo, H., Luo, J., Wang, S., & Zhang, Y. (2020). Trichome regulator SlMIXTA-like directly manipulates primary metabolism in tomato fruit. Plant Biotechnology Journal, 18(2), 354–363. 10.1111/pbi.13202

44. Zhang, S., Chen, H., Guo, S., Wang, C., Jiang, K., Cui, J., & Wang, B. (2024). *Artemisia annua ZFP8L* regulates glandular trichome development. Physiologia Plantarum, 176(4), e14461. 10.1111/ppl.14461

45. Zhao, H., Li, B., Liu, L., Wang, X., Zhu, Z., & Liu, Y. (2025). Deciphering the ″Developmental Code″: A Comprehensive Review of Regulatory Mechanisms in Tomato Trichomes. Journal of Agricultural and Food Chemistry, 73(28), 17358–17373. 10.1021/acs.jafc.5c02904

46. Zheng, F., Cui, L., Li, C., Xie, Q., Ai, G., Wang, J., Yu, H., Wang, T., Zhang, J., Ye, Z., & Yang, C. (2022). Hair interacts with SlZFP8-like to regulate the initiation and elongation of trichomes by modulating *SlZFP6* expression in tomato. Journal of Experimental Botany, 73(1), 228–244. 10.1093/jxb/erab417

47. Zhu, X., Mi, Y., Cao, X., Chen, W., Fan, P., Wang, J., Zhang, Y., Yang, W., Wan, H., Chen, S., Meng, X., Li, J., Shen, S., Huang, M., Zhang, X., Luo, M., Chen, S., Xu, Z., & Sun, W. (2026). A Novel CsYABBY3-CsAS1 Feedback Loop Coordinates Trichome Differentiation and Cannabinoid Biosynthesis in *Cannabis sativa* L. Advanced Science, e75055. 10.1002/advs.75055

